# Mutual anticipation can contribute to self-organization in human crowds

**DOI:** 10.1101/2020.08.09.215178

**Authors:** Hisashi Murakami, Claudio Feliciani, Yuta Nishiyama, Katsuhiro Nishinari

## Abstract

Human crowds provide paradigmatic examples of collective behavior emerging through self-organization. Although the underlying interaction has been considered to obey the distance-dependent law, resembling physical particle systems, recent findings emphasized that pedestrian motions are fundamentally influenced by the anticipated future positions of their neighbors rather than their current positions. Therefore, anticipatory interaction may play a crucial role in collective patterning. However, whether and how individual anticipation functionally benefits the group is not well-understood. We suggest that collective patterning in human crowds is promoted by anticipatory path-seeking behavior resulting in a scale-free movement pattern, called the Lévy walk. In our experiments of lane formation, a striking example of self-organized patterning in human crowds where people moving in opposite directions spontaneously segregate into several unidirectional lanes, we manipulated some pedestrians’ ability to anticipate by having them type on a mobile phone while walking. The manipulation slowed overall walking speeds and delayed the onset of global patterning, and the distracted pedestrians sometimes failed to achieve their usual walking strategy. Moreover, we observed that the delay of global patterning depends on decisions made by pedestrians who were moving toward the distracted ones and had no choice but to take sudden large steps, presumably because of difficulty in anticipating the motions of their counterparts. These results imply that mutual anticipation between pedestrians facilitates efficient transition to emergent patterning in situations where nobody within a crowd is distracted. Our findings may contribute to efficient crowd management and inform future models of self-organizing systems.

## Introduction

Human crowds engage in a rich variety of self-organizing behaviors, as do other animals, such as birds flocking and fish schooling [1–3]. The study of human crowd behavior and modeling pedestrian flows have become crucial research fields to help manage mass events and pedestrian transportation safely and comfortably. Although some collective patterns of organization in human crowds such as stop-and-go waves and crowd turbulence may lead to serious disasters, such as deadly trampling accidents [4–6], there are other phenomena that provide functional benefits to the group even if there is no conductor or external control. Lane formation, where unidirectional lanes are spontaneously formed in bidirectional pedestrian flows, is the most striking example of effective self-organization to reduce the risk of collisions between pedestrians and enhance the efficiency of traffic flow [7–9]. To accomplish efficient crowd management and minimize risks, it is therefore important to understand individual interactions underlying collective human behavior.

Most previous models describe pedestrian dynamics as an interaction potential, or a “social force” inspired from the repulsive potential among physical particles [10–12]. Although these physics-inspired models have significantly improved our understanding of pedestrian crowd dynamics, they do not completely explain the empirical observations. Especially, recent empirical and experimental findings have stressed that inter-individual interaction is essentially anticipatory in nature, rather than a distance-dependent physical force [9, 13–18]. The motions of pedestrians are influenced by anticipated future positions of their neighbors rather than their current positions. This suggests that pedestrians in a crowded neighborhood are not just passively repelled by other pedestrians, but they actively find a passage through a crowd by anticipating and negotiating with neighbors to avoid collisions in advance. Indeed, we previously conducted bidirectional flow experiments regarding lane formation where participants have to deal with people moving toward them and observed that, during formation of lanes, they fluctuate from the desired direction by a Lévy walk process, which is an optimal, scale-free movement strategy to search for unpredictably distributed resources. At the same time, they behave as Brownian walkers in terms of their lateral movements against the desired direction, with minimum fluctuations in the absence of an oncoming crowd when there is no need to anticipate the actions of their walking counterparts [9].

The anticipatory nature in human interactions has been reported in a wide range of pedestrian situations from bottlenecks to hallways, regardless of whether the environment is indoor or outdoor, and seemed to be fundamental for collective human behavior [14]. Understanding the relationship between individual anticipatory behaviors and collective patterning, therefore, is an important issue. Anticipating the motion of neighbors may attenuate head-on collisions between individuals in advance, help to smooth a pedestrian’s pathway through a crowd, and facilitate efficient transition to emergent patterning. However, despite the unprecedented growth of crowd experiment research [19], the influence of anticipatory behaviors on self-organization in human crowds remains poorly understood. In particular, there has been no attempt to experimentally intervene in studies of the cognitive ability of pedestrians’ anticipation.

In this study, we tested the hypothesis that collective patterning in human crowds is facilitated through anticipatory path-seeking behavior. We conducted an intervention experiment of lane formation where some participants were asked to walk while using mobile phones because the use of a mobile phone while walking was thought to interfere with their ability to anticipate. Previous experiments on gait characteristics of single pedestrians reported that mobile phone tasks made pedestrians’ perceptual visual fields narrow when they looked down and focused on the phone, and they distracted visual attention such as scanning frequency around the surrounding environment (e.g., oncoming traffic) [20, 21]. Therefore, distracting pedestrians via a mobile phone task should make it more difficult for them to anticipate their neighbors’ motions because pedestrian motion is strongly influenced by other oncoming pedestrians even when they are well-separated [14]. In addition, it may also be difficult for pedestrians walking toward distracted pedestrians to anticipate the actions of distracted walkers. Thus, self-organization in human crowds facilitated by anticipation may be inferred when this type of distraction causes a deterioration of not only local collision risks but also of global patterning.

We demonstrate that the distraction slows overall walking speed and delays the onset of global patterning. Moreover, the distracted pedestrians had difficulty in achieving their usual walking strategy, the Lévy walk [22–24]. Furthermore, we observed that the delay of global patterning was associated with decisions made by pedestrians moving toward the distracted ones who are forced to take sudden large steps, probably because of difficulty in anticipating the motions of distracted pedestrians. These results are consistent with the concept of self-organization in human crowds facilitated by anticipation. Our findings provide insights into understanding the important role of individual cognitive abilities of anticipation in self-organizing systems.

## Results

### Influence of visual distraction on emergence of lane formation

We conducted experiments where two groups (27 pedestrians each) participated in bidirectional flows in a straight mock corridor (Fig. S1) (see Materials and methods). We imposed an additional task on three participants in one of the two groups to visually distract the pedestrians and disrupt potential anticipatory interactions among them (Fig. 1A). As the distraction, we asked each of these three people to use a mobile phone while walking. We hypothesized that distracted pedestrians located in the front of the group, which could directly collide with the oncoming crowd, would have the most influence on overall crowd dynamics in this bidirectional flow experiment. To test this, we set three experimental conditions (front, middle, and rear) in addition to a baseline condition in which nobody was distracted. In the front condition, three participants in either group were randomly selected to perform the additional task while walking and assigned to the front-most three positions of their respective group. The mid- and rear conditions were the same as the front condition except that the distracted participants were assigned to the middle or rear positions, respectively. The experiments were replicated 12 times under each condition.

**Fig. 1.**
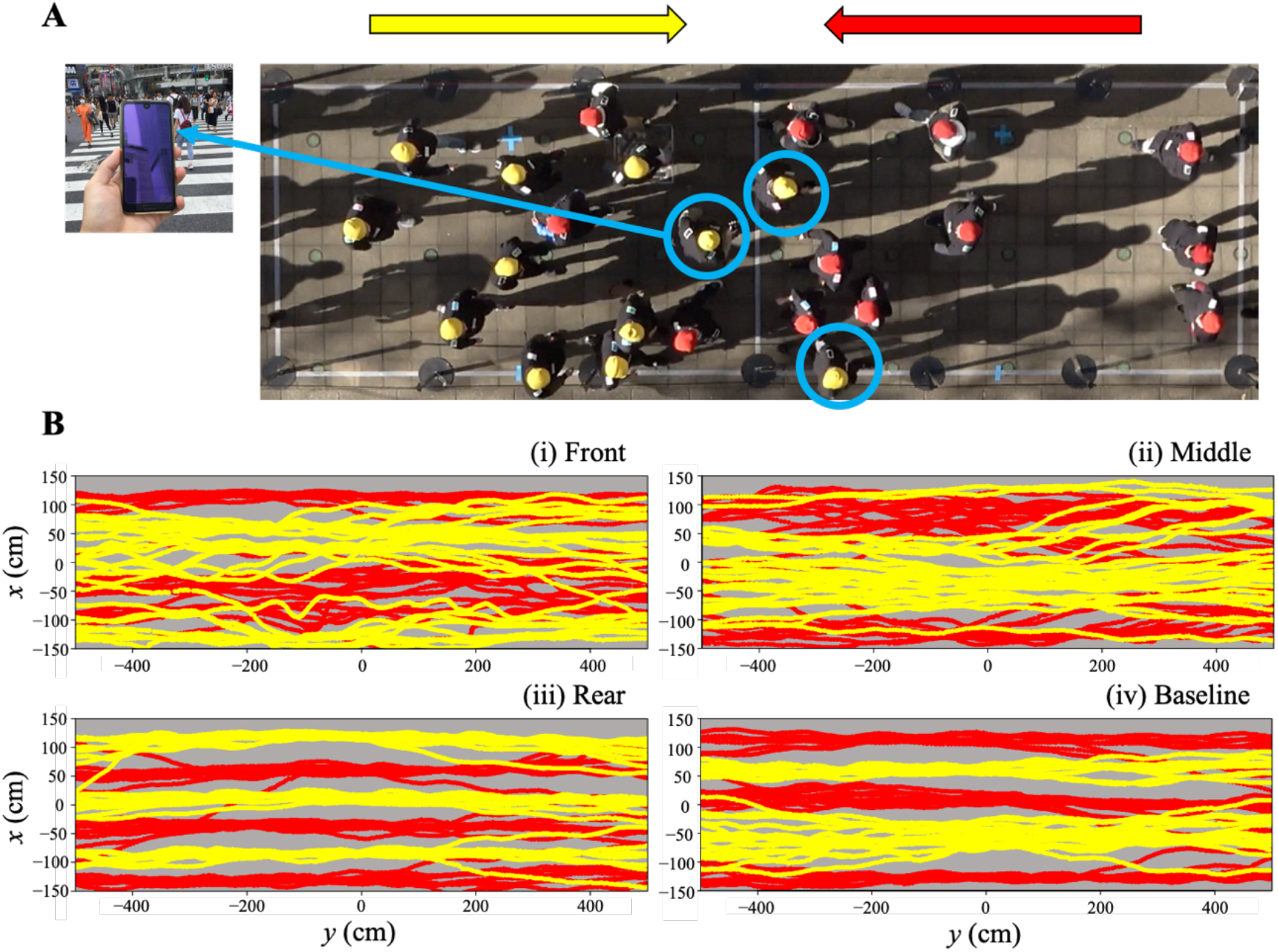
Bidirectional flow experiments with distracted pedestrians. (*A*) Snapshot from an experiment under the front condition (the distracted pedestrians located in the front of a group), where locations of the distracted pedestrians are marked by blue circles. (*B*) Representative examples of reconstructed pedestrian trajectories under (i) front, (ii) middle, (iii) rear, and (iv) baseline conditions. Yellow (red) lines represent pedestrians moving from left (right) to right (left).

Figure 1B shows the development over time of the traffic organization in bidirectional pedestrian flow under each condition. Pedestrians from the two groups initially walked in opposite directions along the corridor, deviating from the direct path to the destination as necessary to seek their passages through a crowd and avoid oncoming pedestrians. Subsequently, they self-organized into two or more unidirectional lanes, and then they followed straight courses along the generated lanes. The distracted pedestrians in the front condition seemed to disturb overall behavior more than they did in the other conditions.

To account for the above observation quantitatively, we first calculated each pedestrian’s walking speed defined as travelled distance in the measurement area divided by crossing time, where the crossing time is the time that a pedestrian took from entering the measurement area to leaving it (Fig. 2A). Pedestrians in the front condition were significantly slower than those in the baseline condition (vs. front: *W* = 259770, *P* < 0.001; vs. middle: *W* = 221949, *P* = 0.06; vs. rear: *W* = 218982, *P* = 0.11), suggesting that distracted pedestrians in the front condition influenced overall flow as we expected.

**Fig. 2.**
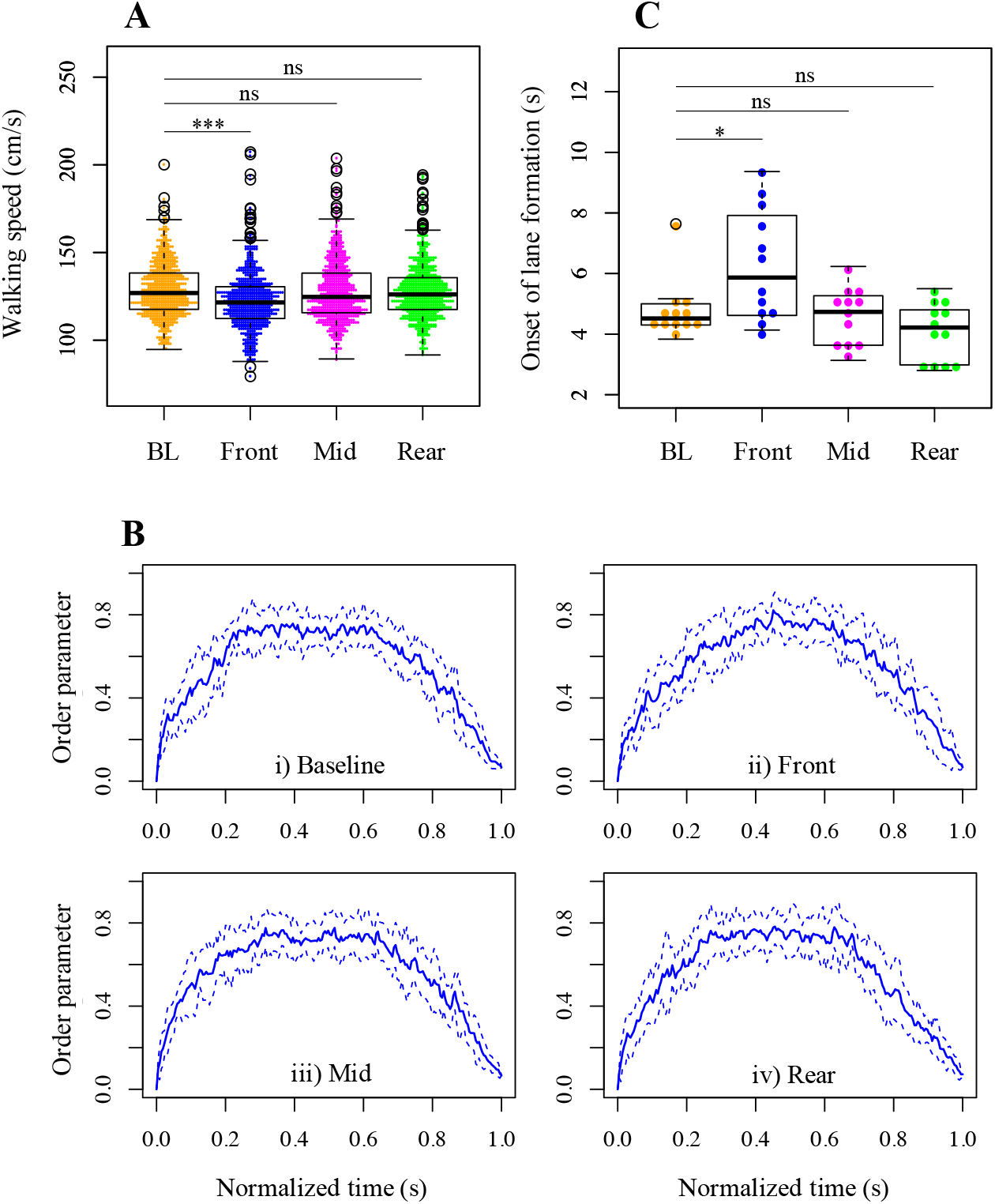
(*A*) Pedestrians’ walking speeds under each condition. Each data point represents a pedestrian. (*B*) Time development of the order parameter under the i) baseline, ii) front, iii) middle, and iv) rear conditions, from the first pedestrian entering the measurement area to the last pedestrian leaving. Time was normalized to be given a relative time axis ranging from 0 to 1. The blue lines show the mean ± standard deviations of the order parameter (solid and dotted lines, respectively). (*C*) Onset of lane formation under each condition. Each data point represents a trial. Asterisks in (*A* and *C*) indicate significance (*** *P* < 0.001; * *P* < 0.05; ns *P* > 0.05). Box-and-whisker plots in (*A* and *C*) represent the median of the data (central thick line), data between the first and third quartiles (box), data within 1.5× the interquartile range of the median (whiskers), and outliers (unfilled circle).

To investigate the self-organization process of lane formation, we calculated the order parameter, which is widely used for measuring the stratification degree of bidirectional flow [8, 9, 25, 26]. The order parameter was defined by discretizing the measurement area into several rows along the corridor and averaging the proportion of individuals moving in the same direction in each row (for details, see text S1). Its value takes 0 when the same number of pedestrians in a row move in different directions or when no individuals occupy a row, and it approaches 1 (which suggests clear lane formation) if most pedestrians in a row move in the same direction. Figure 2B shows plots of the order parameters as a function of time. Under the baseline condition, the plot looks roughly like a trapezoid consisting of 1) a steady increase in the order parameter, which suggests progressive development of lanes; 2) a plateau during a certain period of time, which suggests maintenance of formed lanes; and 3) a steady decrease, which suggests the gradual collapse of lanes. This result is in agreement with our previous study [9]. This trend was also observed in the rear condition and less clearly in the middle condition, but it was not observed in the front condition. Under the front condition in particular, we observed that lanes develop more slowly than under the other conditions. To verify this, we investigated the onset of lane formation by calculating the time at which the order parameter exceeded 0.8 for the first time after the beginning of a trial. This procedure was introduced and justified in [9] (also see text S1). We found that the onset of lane formation in the front condition was significantly later than in the baseline condition but not in the other conditions (vs. front: *W* = 33.5, *P* = 0.04; vs. mid: *W* = 72.0, *P* = 0.76; vs. rear: *W* = 96.5, *P* = 0.92; Fig. 2C). This suggests that only the distracted pedestrians in the front condition disturbed the facilitation of self-organization.

In summary, mobile phone distractions significantly influenced overall walking speeds and the onset of lane formation, especially when the distracted pedestrians were located in the front of the group. The following questions thus arise: how is individual behavior, such as local collision avoidance and the path-searching property of movement in particular, influenced by the distraction, and how is the effect related to global patterning? We answer these questions in the path-seeking section below.

### Local collision avoidance: slow-down/speed-up behaviors

In this section and hereafter, we focused on pedestrians’ behaviors until the onset of lane formation, where path-seeking behavior most likely happens [9], in the baseline condition and front-condition. We assumed that the mobile phone distraction would increase the risk of local collision among pedestrians and that pedestrians would need to perform sudden large turns or steps to avoid collision. In other words, if pedestrians anticipated the motions of oncoming neighbors, they could avoid each other in advance, and therefore they would not take sudden large steps. To verify this assumption, we analyzed pedestrians’ behaviors in more detail through a stratified analysis where we divided the front-condition data into three subgroups: 1) distracted, distracted pedestrians; 2) same-directed, pedestrians walking in the same direction as the distracted ones; and 3) opposite-directed, pedestrians walking in the opposite direction as the distracted ones.

We first investigated the temporal features of pedestrians’ turns. If pedestrians performed collision avoidances locally and suddenly, they would slow down to avoid a collision, make turns with a relatively large turn angle, and then increase their speed to catch up with the preceding pedestrians. As indicators, we determined changes in an individual’s walking speed by calculating differences between two consecutive speed values (i.e., scalar quantities of the velocities) at time *t* and at time *t* – *dt*, where *dt* was the time period and *dt* = 1 (s). We determined angular deviations of the velocity vector from the desired direction (i.e., the direct path to their destination: the *x* axis along the corridor length in this case) by calculating the angle that the velocity vector made with the *x* axis at time *t*. Figure 3 shows the two-dimensional frequency distribution of the indicators with bins of 10 (degrees) × 5 (cm/s). It is clear that the distribution in the baseline condition has a narrower range, with smaller changes in speed and deviation than the others, suggesting that pedestrians under the baseline condition did not perform sudden turns, probably because of anticipation. If we consider the absolute value of the product of the above indicators as an index of a sudden turn, the index in the baseline condition was significantly smaller than the others (see Fig. S2). Moreover, we compared the angular deviation at time *t* with changes in walking speed before a turn calculated from the velocities at time *t* – *dt* and *t* – 2*dt*; we also compared those after a turn calculated from the velocities at time *t* + *dt* and *t*. In these comparisons, we observed that distracted pedestrians and opposite-directed ones tended to slow down before and speed up after large deviations. These results are consistent with our original assumption. Furthermore, although the above analysis focused on temporal features of pedestrians’ behaviors, similar results were obtained when focusing on the spatial features of their behavior (see Fig. S3).

**Fig. 3.**
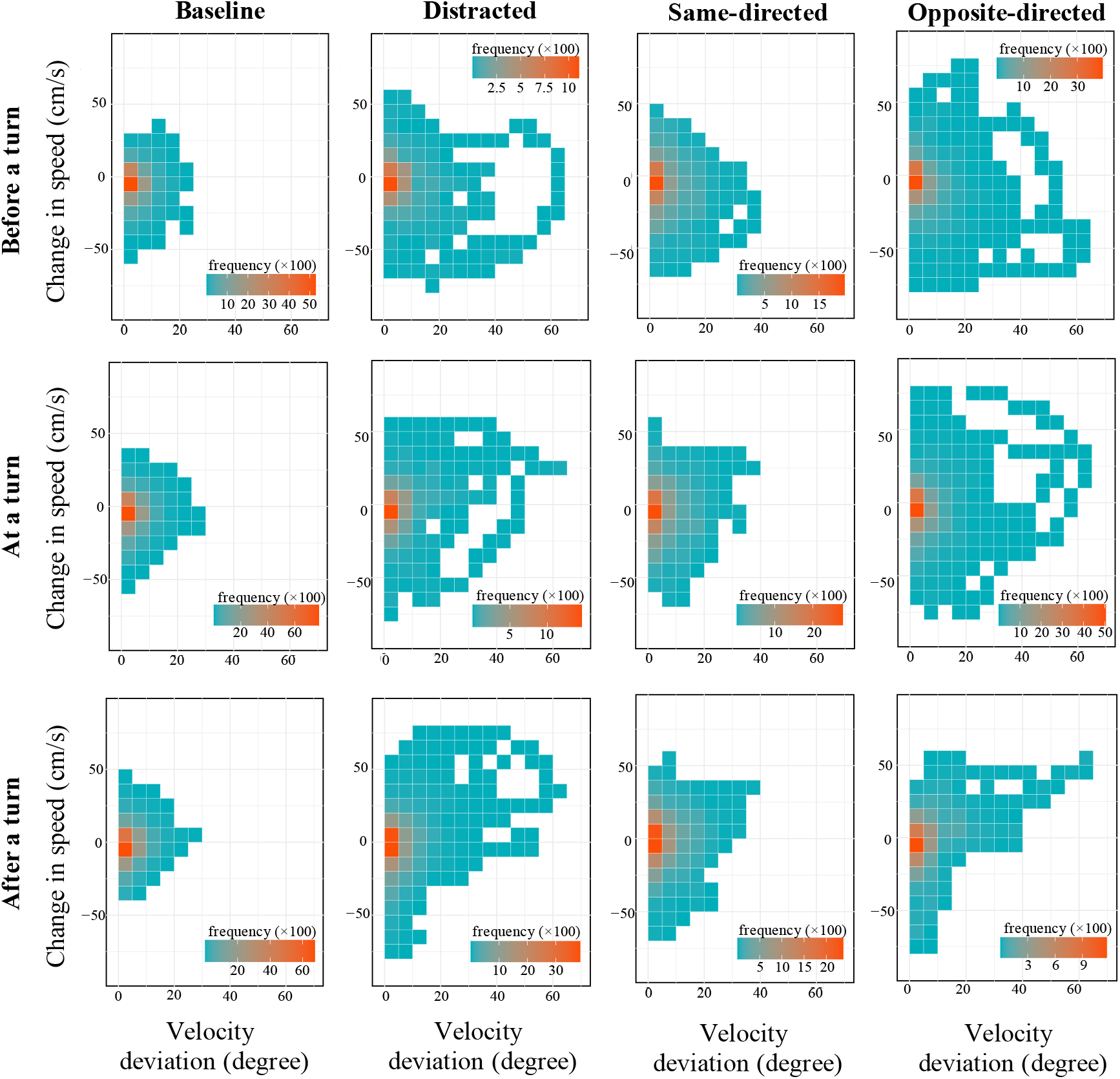
Two-dimensional frequency distributions of change in speed and angular deviation of the baseline, distracted, same-directed, and opposite-directed groups at a turn, before a turn, and after a turn, with bins of 10 (degrees) × 5 (cm/s).

### Path-seeking behavior through a crowd and global patterning

When pedestrians anticipate their neighbors’ motions to seek their paths in dynamic pedestrian flow, this path-seeking behavior involves them deviating from their desired directions, such as in the Lévy walk process [9]. A Lévy walk is a special class of random walks where step length *l* follows a power-law function *P*(*l*) ~ *l*^−*μ*^, where *μ* ∈ (1, 3] represents a power-law exponent; the step length *l* specifies the distance between two consecutive points often marked by a directional change in a walker’s movements. Here we investigate how a distraction influenced pedestrians’ path-seeking behavior and discuss the relationship between the path-seeking behavior and global patterning. To this end, we decomposed the individual trajectories into an *x*-component (along the corridor length) and a *y*-component (perpendicular to the corridor length), focusing on the latter (i.e., the lateral movements of pedestrians), which is considered to be the most related to searching activities. Here, step lengths were calculated as the *y*-components of the distances between consecutive turning points, which were defined as direction changes in the *y*-component of an individual’s trajectory. We tested whether pedestrian movements followed the truncated power law (Lévy walking) as compared with an exponential distribution (Brownian walking). To do the comparison, we calculated Kolmogorov–Smirnov (KS) statistics to determine the goodness-of-model fits (GOF) selected by the Akaike Information Criterion (AIC) with maximum likelihood estimations (for details, see text S1) [27–34].

Figure 4A–D shows the step-length rank-distributions for each trial (see Figs. S4–7 for plots with the best-fitting truncated power-law and exponential models). The plots of the distracted pedestrians trended more upward or downward in each trial than those of the others. For the distracted pedestrians, the statistical results showed that the proportion of trials showing step-length distributions judged to follow the truncated power law out of the total number of trials (12) was the lowest of all groups: 12/12 in the baseline group, 7/12 in the distracted group, 11/12 in the same-directed group, and 12/12 in the opposite-directed group (*μ* = 1.86 ± 0.108, 1.74 ± 0.408, 1.74 ± 0.256, and 1.88 ± 0.133 [mean ± sd], respectively). The tendency of a worse fit to the truncated power law for distracted pedestrians was additionally confirmed through detailed analyses of the differences from the baseline in the AIC weights and GOF (Fig. S8). Moreover, we analyzed the diffusive properties along the *y*-axis using the mean-squared displacement (see text S2). According to this analysis, the diffusive characteristics of distracted pedestrians’ movements varied by trial, even when including the normal diffusive properties that indicate almost straight paths along the corridor with only small lateral deviations. The other subgroups and baseline group demonstrated super-diffusive characteristics of movement patterns in all trials (Fig. S9).

**Fig. 4.**
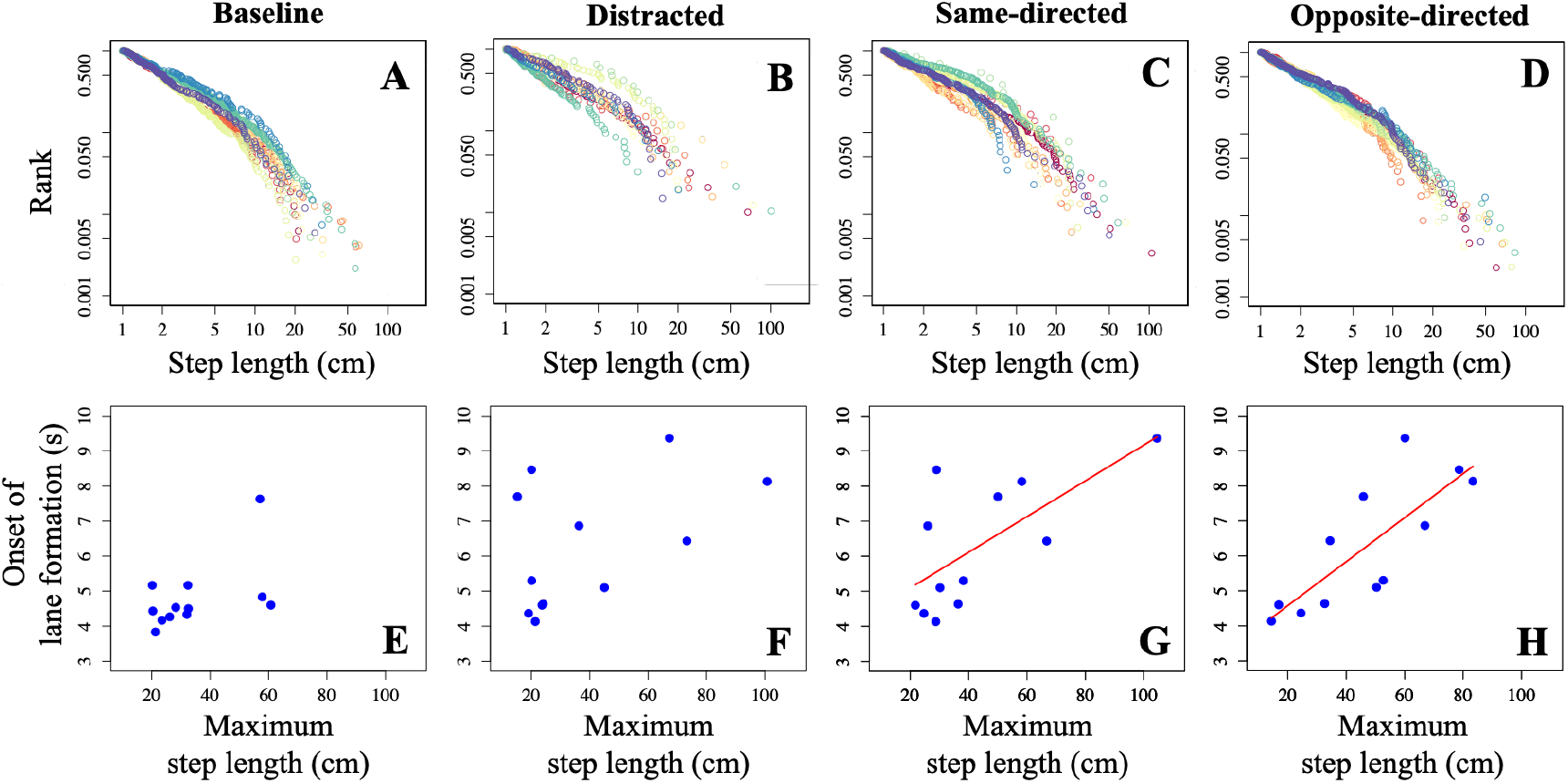
Rank distributions of step length along the *y*-axis in each trial of (*A*) baseline, (*B*) distracted, (*C*) same-directed, and (*D*) opposite-directed groups. Different colors represent different trials. Onset of lane formation as a function of maximum step length of (*E*) baseline, (*F*) distracted, (*G*) same-directed, and (*H*) opposite-directed groups. Red lines indicate fitted linear regressions showing significant effects (*P* < 0.05).

Finally, we investigated the relationship between global patterning and local individual movements, focusing on maximum step length, which represents the order of magnitude of the step-length distribution and implies the level of local collision risk that pedestrians were forced to avoid. As a parameter of global patterning, we employed the onset of lane formation again. Interestingly, we observed that the onset of lane formation was positively affected by the maximum step length of opposite-directed members as well as same-directed ones, whereas there were no significant effects for baseline and distracted pedestrians (baseline: *R*^*2*^ = 0.276, *F* = 3.813, *P* = 0.08; distracted: *R*^*2*^ = 0.238, *F* = 3.131, *P* = 0.11; same-directed: *R*^*2*^ = 0.4541, *F* = 8.32, *P* = 0.02; opposite-directed: *R*^*2*^ = 0.6286, *F* = 16.92, *P* = 0.006; Fig. 4E–H). This suggests a possibility that the delayed global patterning was caused by the decision-making of non-distracted pedestrians, especially opposite-directed members, rather than by that of the distracted pedestrians per se.

## Discussion

In the present study, we addressed whether self-organization in human crowds can be promoted by anticipatory interaction among individuals. To this end, we conducted bidirectional flow experiments, where some participants were asked to walk while typing into mobile phones. Distracting pedestrians at the front of a group resulted in slowdown of overall crowd movement and delayed onset of global lane formation. In addition, the distracted pedestrians sometimes did not perform exploratory walking in the crowd, as observed by their poor path-seeking behavior, even though such exploration is considered to be common in bidirectional lane formation situations [9]. Moreover, the increase of local collision avoidance and the delayed onset of global patterning predicted by maximum step lengths demonstrated the indirect effects of the experimental manipulation on non-distracted pedestrians as well as the direct effects on distracted ones. The distraction thus locally reduced individual anticipation ability and globally disturbed the self-organization process through the propagation of reduced levels of anticipation in the crowd. These results are consistent with the idea that self-organization in human crowds is facilitated by anticipation.

The mobile phone distraction altered behaviors of both distracted and non-distracted pedestrians. Indeed, we observed that distracted pedestrians intensely performed local collision avoidance behavior as compared to normal pedestrians in the baseline condition and sometimes failed to achieve the usual walking strategy. This is consistent with the assumption that pedestrians performing path-seeking behavior, represented by the Lévy walk, can avoid collisions in advance. According to a previous study, distracted pedestrians fixated their eyes approximately 75% of the time on the mobile phone while typing [20]. Although the previous study used only single pedestrians walking through a crosswalk, it would not be surprising if similarly low frequency of scanning surroundings occurred in our experiments. Therefore, it is plausible enough that the distracted pedestrians performed emergency avoidance movements because they were often unaware of their neighbors until immediately before potentially running into them.

Somewhat surprisingly, pedestrians moving toward distracted ones also intensely performed local collision avoidance behavior as compared to pedestrians under the baseline condition, despite showing path-seeking behavior that results in the Lévy walk. This suggests that their anticipatory behavior toward oncoming pedestrians sometimes did not work well, even though they were able to dedicate their visual resources to their walking task. We speculate that this local collision avoidance of opposite-directed pedestrians was caused by their uncertainty of the motions of distracted walkers for the following reasons. First, the opposite-directed pedestrians most likely obtained less information from the gaze direction of the distracted ones, from which humans usually infer other people’s movement trajectories [35]. Because distracted pedestrians looked down and focused their eyes on the phone, opposite-directed ones would be uncertain about the motions of distracted ones. Second, in behavioral neuroscience studies, human decision-making has been considered in a serial “decision-then-action” framework where sensory evidence is collected and then action is taken. However, recent studies have stressed a “decision-during-action” framework because the decision-making may involve deadlines and time constraints for action [18, 36–38]. In this framework, action plans can be prepared and even launched before information from the sensory cues is completely accumulated. In other words, decisions can be made using real-time exploration for sensory cues during action while the information remains uncertain.

We applied these ideas to the local collision avoidance behaviors we observed. In the baseline situation with no particular problems, pedestrians moving toward a destination should balance the benefits of making the correct decision (e.g., safely avoiding collisions) with its costs (e.g., the time constraint to reach the destination), and react accordingly. However, if uncertainty of motion is introduced in the form of oncoming distracted people, pedestrians will have to continue to seek rare information. Consequently, they will sometimes reach the situation where a collision is imminent and turn immediately before colliding with the distracted walkers. In this way, we consider that the distraction in our experiment not only directly deprived the anticipatory ability of the distracted walkers but also indirectly influenced that of the opposite-directed ones. This then induced collision avoidance behavior even though the opposite-directed walkers were showing path-seeking behavior. In other words, collisions can be avoided in advance only if anticipation is performed mutually among pedestrians. Therefore, mutual anticipation may contribute more to a smooth walk than one-way anticipation.

In addition, we observed a link between individual anticipation and the self-organization process in a crowd. The fact that the onset of lane formation under the front condition ranged more widely as compared to the other conditions (Fig. 2C) suggests that the mobile phone distraction had a range of influence, from heavy to moderate. Moreover, the onset of lane formation was statistically predicted by the maximum step length (Fig. 4G, H), suggesting that the delay of global patterning was caused not only by the decision-making of distracted pedestrians, but also by that of the opposite-directed walkers, who were having difficulty in mutually anticipating motions with the distracted ones. These results imply that, if the non-distracted pedestrians can successfully anticipate motions of the distracted people and walk smoothly, collective patterning is achieved rapidly; otherwise, patterning is delayed.

Considering all of the walking strategy results together, because non-distracted pedestrians maintained reasonable path-seeking behavior that appeared as Lévy walk, such a process might guarantee emergent patterning in human crowds. The Lévy walk was originally hypothesized as an efficient foraging strategy for animals with low cognitive ability when searching for unpredictably distributed resources. However, recent findings strongly suggest that the Lévy walk and scale-free supper-diffusive movement patterns play a crucial role even in human mobility, including foraging behavior as well as daily movements in urban areas [28, 39–42]. Our results provide an example of the Lévy walk in human behavior related to self-organization. Although the Lévy walk was hypothesized to have evolved through natural selection because of its searching efficiency, recent findings suggest that it might also be the result of other mechanisms, especially through interactions with a complex environment, including the distribution of conspecifics [22, 29, 43–45]. For example, mussels use it during the formation of spatially patterned beds [29]. This pattern helps to improve individual fitness through complex feedback while individuals interact with an “environment” (i.e., their own distribution). Our results demonstrated that the Lévy walk resulted from interactions in human crowds. Thus, it is likely an important element in the formation of collective patterning in widespread biological systems.

Understanding anticipatory interaction between pedestrians would provide fundamental insights into crowd dynamics and a crucial step toward effective and safe management of mass events. Our results describe how people navigate in close surroundings and the degree of effect distracted people have on a crowd. Potential real-life applications in daily crowd management include efficient signage, customized navigation, and wayfinding. Using an eye-tracking system would be an effective measure to fully explore the effects of various degrees of anticipation on interactions among pedestrians according to the difficulty of the distraction task and the number of distracted pedestrians (i.e., their proportion to the total). Moreover, we experimentally investigated the functional anticipatory interactions in human crowds, but it is important to explore representations in theoretical modeling. We expect that our results will provide quantitative support to existing models, especially vision-based or velocity-based ones [13, 14, 16], some of which are inspired from a cognitive science approach rather than physics-inspired repulsive interactions. In these models, pedestrians *actively* select the path with the longest distance to the predicted collision with the neighbors. Yet, at the same time, the present findings additionally require that pedestrians alter their behavior depending on the uncertainty of oncoming neighbors’ motions. Human crowd models that better incorporate anticipatory behavior will provide a more reliable description and prediction of pedestrian flows and therefore open the way for the improvement of architecture and exit routes, as well as management strategies during mass events. Moreover, future studies focusing on anticipation in human crowds might open new perspectives in other research fields concerned with self-organizing systems in general. For example, anticipatory behavior contributes to cooperative human decision-making through synchronization between two persons [15, 18]; animal groups are considered to achieve robust collective behavior through mutual anticipation [46–50]; and swarm robotics has problems in navigation and collision avoidance in the real world and needs a new approach in industrial applications [51]. Anticipation likely plays a key role in a wide range of self-organizing systems. Thus, we expect that the present findings will become a foothold to build future models of collective human behavior, biological collectives, and swarm robotics.

## Materials and methods

We conducted controlled experiments to study the dynamics of lane formation and its relationship with anticipatory behavior of individual pedestrians, adapting our previous experimental setup and procedure [8,9] (see text S1). For the purpose of simulating bidirectional pedestrian flow at a crosswalk, we employed an experimental, straight mock corridor, mainly composed of a measurement area (10 × 3 m) and two waiting areas (12 × 3 m), one located at each side of the measurement area (Fig. S1). We employed 54 participants divided into two groups of 27 to make balanced bidirectional flows. At the beginning of a trial, each group was in a separate waiting area that had nine predetermined starting lines perpendicular to the long axis of the corridor. Each line was separated by an equal distance (1.5 m). Participants in each group were assigned positions at random, but there were three participants per starting line. At the start signal, they were asked to walk in their usual way toward an exit at the opposite end of the mock corridor.

To visually distract pedestrians and disrupt potential anticipatory interactions among them, we imposed an additional task on three participants in one of the two groups (text S1 and Fig. S10). Inspired by previous studies of mobile phone distraction [20], we provided each of these three people with a mobile phone and asked them to try to solve as many simple arithmetic problems (one-digit addition) as possible while walking. We assumed that distracted pedestrians located in the front of the group, who would be the first to meet or collide with the oncoming pedestrians (i.e., the “crowd”), would have the most influence on overall crowd dynamics in this bidirectional flow experiment. We set therefore three experimental conditions in addition to a baseline condition in which nobody was distracted: front, middle (mid), and rear. In the front condition, three participants in either group were randomly selected to perform the additional task while walking and assigned to the front-most three positions (i.e., behind the first starting line) of their respective group. The mid- and rear conditions were the same as front condition except that the distracted participants were assigned to the middle (the fifth starting line) or rear (the ninth line) positions, respectively. The experiments were replicated 12 times under each condition.

To control for the possibility that a decrease of walking speed induced by the mobile phone distraction [20] might influence the lane formation process rather than anticipatory ability, we conducted additional experiments that were similar to the above ones, but the three selected pedestrians were only asked to walk slowly instead of performing a task on a mobile phone. We confirmed that slow walking per se did not impact lane formation (see text S3 and Fig. S11).

Additional details about the experimental setup, procedures, and analyses are described in text S1. The experiments were conducted at the University of Tokyo in December 2019. This study was approved by the Ethics Committee of the University of Tokyo, and informed consent was obtained from all participants.

## Competing interests

We have no competing interests.

## Authors’ contributions

H.M., C.F., and Y.N. designed the study; H.M., C.F., and Y.N. performed the experiments; H.M. analyzed the data; K.N. provided funding acquisition, project administration, and resources; and H.M. wrote the paper.

## Acknowledgements

We thank all of the participants and members of Nishinari laboratory for their help during the experiments. We are also grateful to Dr. Daichi Yanagisawa, Dr. Yuki Oyama, and Dr. Masahiro Furukawa for their helpful discussions. This work was supported by JSPS KAKENHI grant numbers JP20K20143, JP20K14992, and JP19K20384, and JST-Mirai Program grant numbers JPMJMI17D4 and JPMJMI20D1.

## Notes

### Competing Interest Statement

The authors have declared no competing interest.

