## Supplemental information for "Mutual anticipation can contribute to self-organization in human crowds"

**Supporting information for:**
### Mutual anticipation can contribute to self-organization in human crowds

Hisashi Murakami[1]*, Claudio Feliciani[1], Yuta Nishiyama[2], Katsuhiro Nishinari[1,3]

[1]Research Center for Advanced Science and Technology, The University of Tokyo, Meguro-ku, Tokyo, Japan
[2]Information and Management Systems Engineering, Nagaoka University of Technology, Nagaoka, Niigata, Japan
[3]Department of Aeronautics and Astronautics, Graduate School of Engineering, The University of Tokyo, Bunkyo-ku, Tokyo, Japan

*Corresponding author:
H. Murakami, (HM)

**This SI contains text S1 to S3, Figures S1 to S11, and references for the SI.**

### Supplementary Information

#### Text S1. Materials and methods

**Experimental setup and procedure** The experimental setup was adapted from that described in our previous studies [1, 2]. Fifty-four male participants were recruited for the study from university students (mean age = 21.0 ± 2.3 years). We employed a straight corridor composed of three main parts: a measurement area (10 × 3 m) centered in a corridor and two waiting areas (12 × 3 m), one on each side of the measurement area (Fig. S1). Buffer zones (2 × 3 m) between the measurement and waiting areas allowed participants to stabilize their walking speed when entering the measurement area. Each waiting area was divided into sections along its length with nine lines perpendicular to the long axis of the corridor. The lines were separated by an equal distance of 1.5 m and were used as starting lines in the experiments. Destination areas were set at the end of each waiting area.

The experimental procedure was similar to that in our previous experiments [1, 2]; however, we revised some details to intervene in the anticipatory interactions among pedestrians in crowds by making some of the participants distracted by means of mobile phone use. We first divided participants into two groups of 27 to have balanced bidirectional flows. Then, participants in each group were instructed to take random positions on randomly selected starting lines in each waiting area, with three participants per line. At the start signal, they were asked to walk in their usual way toward the exit at the opposite end of the mock corridor and to continue walking even after exiting the measurement area until entering the destination area on the opposite side to avoid the measurement area becoming jammed at the end of the experiment. Before starting the experiments, we informed all participants that there could be three people who would walk while using a mobile phone. We did this to decrease the risk of direct collision involving physical contact that could be induced by the distraction and to ensure that the same number of participants knew of the existence of distracted pedestrians during the experiment (i.e., to avoid a sort of progressive learning process in which participants gradually begin to understand details of the experimental procedure).

Three participants were randomly selected from one of the groups to use a mobile phone during the experiments to distract pedestrians visually and intervene in the anticipatory interactions among them. The three phone users were placed in only one of the two groups during a trial to decrease the risk of collision among distracted pedestrians, and we did not observe any hard physical contact in any of our experiments. Each distracted participant performed the mobile phone tasks for as many as two replicates. The three distracted participants were instructed on how to perform the mobile phone tasks immediately before each trial as described below.

In addition to observing the distracted pedestrians' behavior, we attempted to observe the behavior of pedestrians who walked toward the distracted pedestrians (see the main text). We hypothesized that distracted pedestrians located in the front of the group, which could directly collide with the oncoming crowd, would have the most influence on overall crowd dynamics in our bidirectional flow experiment. To test this, we considered four positions (front, middle, rear, and baseline) as described in the main text.

We conducted two pretest trials without the mobile phone distractions to confirm participants had correctly understood instructions given them before conducting the main experiments. The main experiments were divided into two sessions of six blocks each. As mentioned previously, we divided participants into two groups that were maintained during a single session, but the group members were randomly shuffled between the two sessions. In each block, four trials with four different conditions (i.e., front, middle, rear, and baseline) were conducted in a randomized order. In total, experiments were replicated 12 times under each condition.

In all experiments, we recorded the measurement area from above using a Sony FDR-AX60 camcorder (3840 × 2160 pixels, 30 frames/second) fixed at a height of 21 m. The participants wore black T shirts and different colored caps (yellow and red) to make it easier to distinguish the different flows from one another. From the video images, we tracked the time series of the individuals' positions frame-by-frame, using image-processing software (PeTrack [3]).

**Mobile phone task** Distracted walking has become a major cause of pedestrian accidents so that many researchers have begun to investigate the influence of mobile phone distraction, especially on the behavior of a single pedestrian at a crossing. To introduce a mobile phone distraction in a crowd experiment, we selected a typing task, which has been shown to clearly distract pedestrians' visual attention in terms of scanning frequency, fixation points, and fixation times toward traffic environments [4, 5]. Inspired by previous studies, we employed a simple arithmetic task with a single-digit addition problem similar to Kraepelin's arithmetic test. A simple interface was constructed with Android Studio version 3.5 (https://developer.android.com), where a single-digit addition problem, numeric keyboard, and OK button were presented on the display (Fig. S10). After the user types in an answer and pushes the OK button, the next problem is presented regardless of whether the answer was correct or incorrect. To remove the influence of different mobile phones, the same phone model was used by all participants in the distracted role (UMIDIGI A3, 5.7-inch display, aspect ratio 19:9, 186 g in weight).

We asked participants to start solving problems from the moment they heard the start signal until they entered the destination area and to endeavor to solve as many problems as possible. The number of answers and the proportion of correct answers were recorded for each participant. After one rear-condition trial, however, one mobile phone had a technical problem, so the number of answers and proportion of correct answers were lost for that one sample (i.e., we obtained 35 samples for the rear condition and 36 samples for the other conditions). We then checked if there were any differences in performance of calculations among the conditions. To control for the effect of differences in walking distance and speed among conditions on the number of answers, we estimated the number of answers per second by using the number of answers, distance between the start lines and the start of opposite destination area (i.e., the approximate distance that pedestrians walked during the calculation period), and the median value of the walking speed (travelled distance in the measurement area divided by the crossing time) of the distracted pedestrians under each condition. We confirmed that there were no significant differences among the conditions (i.e., front, middle and rear; see Fig. S10) in number of answers per second (Kruskal–Wallis test: $H = 2.83$, $P = 0.24$) or the proportion of correct answers (Kruskal–Wallis test: $H = 1.34$, $P = 0.51$) under each condition.

**Definition of the order parameter and onset of lane formation** To investigate the self-organizing process of lane formation, we calculated the order parameter, which was originally used for colloidal fluids, but is now used for measuring the stratification degree of various bidirectional flows. The order parameter was measured by discretizing the measurement area into 0.2 × 0.2 m cells (50 columns × 15 rows), as proposed in [1] and justified in [6]. We then focused on the rows of aligned cells along the corridor. The local order parameter $\varphi_j$ in the $j$-th row was calculated as:

$$\varphi_j = ((n_j^L - n_j^R) / (n_j^L + n_j^R))^2 \text{ if at least one pedestrian occupies the row, } 0 \text{ otherwise.} \quad (1)$$

Here, $n_j^L$ and $n_j^R$ are the numbers of left and right walkers in the $j$-th row, respectively. The global order parameter $\Phi$ was then calculated as:

$$\Phi = (1 / N)\, \Sigma^N \varphi_j, \quad (2)$$

where $N$ is the number of rows. $\Phi$ was smoothed over a time window of three frames. Its value becomes 0 when the pedestrians in a row move in different directions or when no individuals occupy the row and approaches 1 if most of the pedestrians in the row move in the same direction, suggesting clear lane formation.

By using the order parameter, we considered the onset of unidirectional lanes as the time at which $\Phi$ first exceeded 0.8, as verified in [2]. We considered this value reasonable in [2] because 1) the approximate time at which $\Phi$ reaches the first (left) corner of the trapezoid shape appears in the order parameter vs. time curve (see Fig. 2B in the main text) appropriately indicates when the pedestrian behavior divides, provided that the data before and after lane formation are reasonably separated; 2) the parameter value does not significantly affect the onset of lane formation as long as it lies in a certain interval; and 3) $\Phi$ reached this value in all trials (this was also true in the present experiment), so the onset of lane formation was determinable in all trials.

**Model fitting** We calculated the exponents and coefficients of the truncated power-law and exponential models that were the best fit for our step-length data. The analysis was based on maximum likelihood estimation and assessed by the Akaike Information Criterion (AIC); the significance of the best-fit models was determined with Kolmogorov-Smirnov (KS) statistics [7–16]. Because the maximum step length of humans and other animals is limited by various conditions such as physiology and spatial constraints, the truncated power law is generally regarded as more biologically plausible than the pure power-law model [7, 8]. The truncated power law is described by the following probability density function of step length $l$:

$$f(l) = (\mu - 1)\,(l_{min}{}^{1-\mu} - l_{max}{}^{1-\mu})^{-1}\, l^{-\mu}, \tag{3}$$

where $\mu$ is the power-law exponent, $l_{min}$ is the start of the data, and $l_{max}$ is the maximum value of the data for the model. The probability density function of the exponential model is

$$f(l) = \lambda \exp[-\lambda(l - l_{min})], \tag{4}$$

where $\lambda$ is the exponent for the model. In previous studies, $l_{min}$ was determined by an iterative procedure [14]. We used 1 cm as $l_{min}$ rather than the iterative procedure described in some previous works to determine $l_{min}$. This procedure is more conservative and less likely to force data into a power-law distribution [8, 16]. $l_{max}$ was set to the maximum observed value. First, the best-fit exponents of the truncated power-law and exponential models were determined by maximum likelihood estimation (MLE). Second, the log-likelihoods of the best-fit truncated power-law and exponential models were calculated. Third, the AIC weights were calculated from the log-likelihoods of both models; the larger the AIC weight was, the better the model support for an observed step-length distribution. Finally, the goodness-of-fits (GOFs) of both models were calculated by using the KS statistics with an iterative procedure [8, 14], where 1000 simulated datasets were generated using the calculated exponent from the sample for the truncated power-law and exponential models. Each of these simulated datasets was then fit by using MLE procedures. A KS statistic was calculated for each of these datasets. The proportion of datasets with larger KS statistics than the empirical dataset was the *P* value. The reliabilities of the models were then assessed from the resulting *P* values: the smaller the *P* value, the less plausible the model fit [8, 14]. We considered that the truncated power-law model was plausible for the data if the AIC weight of the truncated power law was larger than that of the exponential, the *P* value of the GOF of the truncated power law was greater than 0.1, and the *P* value of the GOF of the exponential was smaller than 0.05.

**Statistical analysis in the main text** One-tailed Mann–Whitney–Wilcoxon tests were used to test differences of walking speed and the onset of lane formation between the baseline and each other condition (front, middle, and rear). Linear regression analyses were used to examine the effects of the maximum step length on the onset of lane formation. When making multiple comparisons, *P* values were adjusted by using false discovery rate control [17]. All statistical analyses were conducted using R version 3.6.0 (The R Foundation for Statistical Computing, Vienna, Austria).

**Text S2. Mean-squared displacement**

To characterize the diffusiveness of pedestrian movements, particularly the lateral movements of pedestrians (i.e., deviation from the desired direction, from the *x*-axis in this case), we calculated the mean-squared displacement (MSD) for *y*-components (perpendicular to the corridor length) of pedestrian trajectories before lane formation, which is where path-seeking behavior most likely occurs [2]. The MSD of the *y*-component was computed as a function of time: $\delta y^2(t) = \langle|y(s + t) - y(s)|^2\rangle$, where $y(s)$ denotes a pedestrian's position along the *y* axis at time *s*, and the triangular brackets indicate averaging over the sample trajectories over all times *s*. Therefore, $\delta y^2(t)$

measures the average distance along the $y$ axis during time $t$. Most natural diffusion processes are well described by the power law: $\delta y^2(t) \sim t^{\alpha}$, where the diffusion exponent $\alpha$ ranges from 0 to 2. Brownian (normal) diffusion is characterized by $\alpha = 1$. If $\alpha > 1$, the diffusion is faster than Brownian diffusion and is referred to as super diffusion (the extreme case with $\alpha = 2$ is called ballistic diffusion).

We observed that, at later times, the slopes trended upward or downward or undulated for each condition, in accordance with the results of a previous study [2]. This can be explained by the relatively low sample number at later times and/or the higher probability of encountering the corridor border by pedestrians with relatively longer movements. To determine the inflection point where diffusive regimes change, we fitted a two-segment piecewise regression to logarithmically transformed squared displacement vs. time plots [8] for each trial of the baseline and each subgroup. We iteratively split each dataset into two subsets: the first $i$ points in time and the last $n - i$ points, where $i$ was allowed to iterate from 3 to $n - 3$. Ordinary least squares regressions for log(squared displacement) regressed against log(elapsed time) were then fit to the two subsets in each iteration. GOF was calculated as the sum of the squared residuals from the two segments of the piecewise regression, and the value of $i$ that minimized the sum of the squared residuals was used to identify the best fit model. We then considered only the primary segment and its slope that resulted from the above best fit procedure. Figure S9 plots the mean-squared displacements before lane-formation of each trial of the baseline and each subgroup of the front condition. We observed that the diffusive characteristics of distracted pedestrian movements tended to be more widely varied among trials that included normal diffusions with an $\alpha$ close to 1, whereas the others showed super-diffusive characteristics of movement patterns in all trials.

**Text S3. Additional experiments with slow-walking pedestrians**

To verify the possibility that the decrease of walking speed induced by mobile phone distraction influenced the lane formation process rather than anticipatory ability, we conducted additional experiments. The slow-walking experiments were conducted on the same day as the other experiments. The experimental procedure was same as the main experiment except for one instruction; the three selected pedestrians were asked to walk slowly instead of performing mobile phone tasks. As with the main experiment, we considered the same three conditions regarding where these three slow-walking pedestrians were located in the group: front, middle, and rear. The experiments were replicated three times under each condition.

As a result, we first observed that overall walking speed under the front condition was less than that of the baseline, which was similar to the main results (Fig. S11A). However, overall walking speed under the front condition in this additional experiment was less than that in the main experiment (90.2% and 95.7% less than baseline in terms of median value, respectively). Nevertheless, instructing pedestrians to walk slowly did not impact the onset of the lane formation (Fig. S11B). Moreover, we did not observe the intense collision avoidance behavior that was observed in the main experiments (Fig. S11C, D) (also see the main text). We, therefore, considered that the slow walking did not have an influence on the formation of lanes.

Figure S1

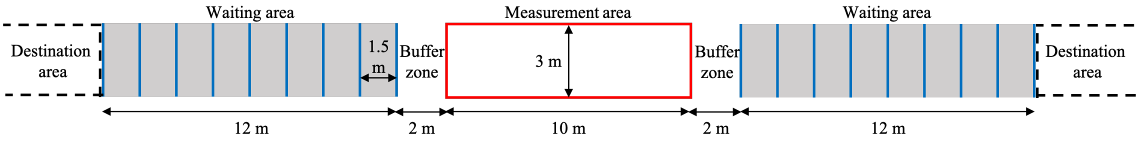


**Fig. S1**. Illustration of the experimental corridor. The corridor includes a measurement area sandwiched between two waiting areas, with buffer zones between the measurement and waiting areas. The buffer zones allow participants to stabilize their walking speed when entering the measurement area from the waiting area. Each waiting area includes nine lines used as starting locations (vertical blue lines). Destination areas were set at each end, outside of the waiting areas.

Figure S2

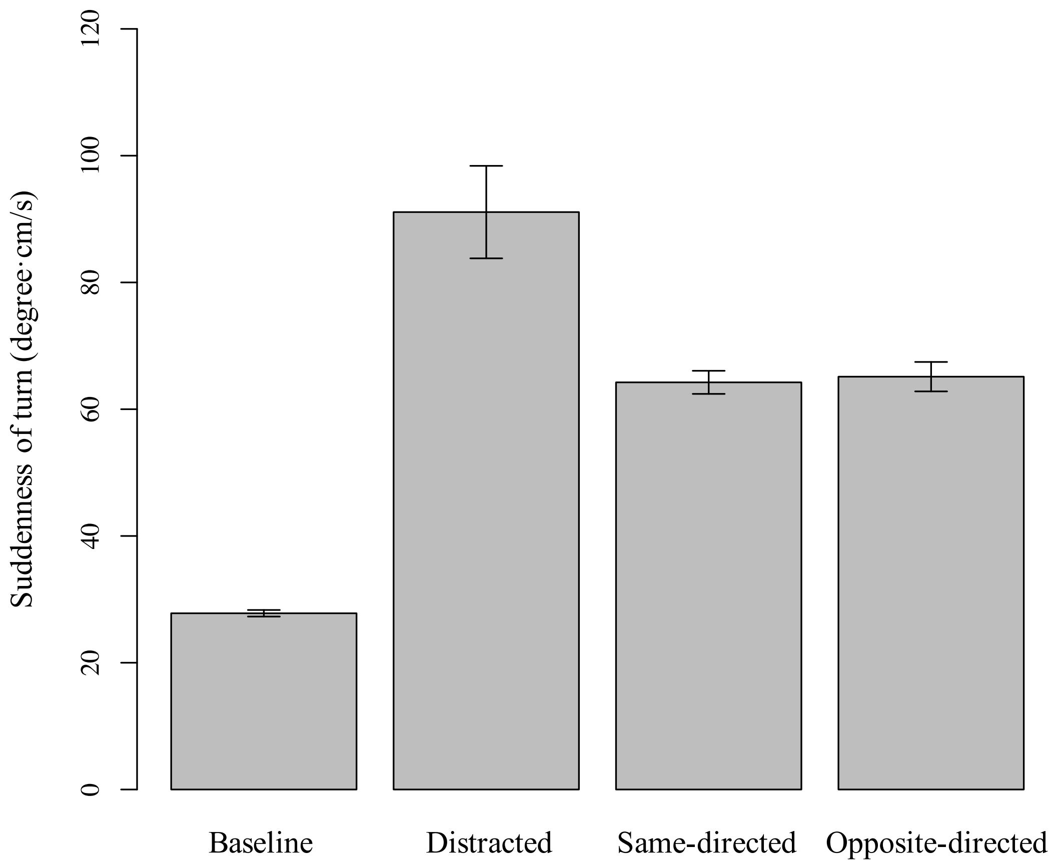


**Fig. S2.** Suddenness of turns, calculated as the absolute value of the product of velocity deviation and change in speed. Error bars are 95% confidence intervals. When compared to the baseline, the values of the other conditions are far larger, suggesting that pedestrians under the baseline condition did not perform sudden turns, most likely because of their ability to anticipate, whereas participants in the other conditions were forced to perform them.

Figure S3

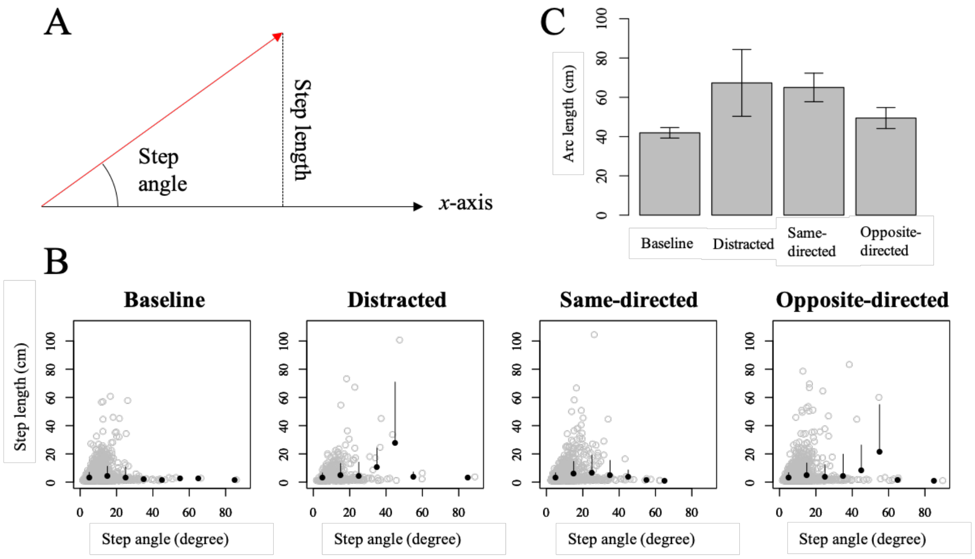


**Fig. S3.** Analysis of local collision avoidance behavior focusing on the spatial features of pedestrian behavior by means of step length and angle. (*A*) Illustration of step length and angle. The red arrow represents a single-move step (the distance between two consecutive turning points defined as direction changes in the *y*-component of an individual's trajectory, as in the main text). Here we considered the step length as the *y*-component of the distance and the step angle as an absolute value of the angle deviation of the step from the desired direction (*x*-axis in this case). Considering the results of our analysis of the temporal features of pedestrian behavior (see the main text), we assumed that these two indicators would take large values at one point in time if pedestrians performed sudden large steps. (*B*) Plots of step length against step angle. Black filled circles (error bar: standard deviation) are the binned scattered data (gray circles), with an interval of 10 degrees. The distracted and opposite-directed groups demonstrate some steps with large lengths and angles as compared to those of the baseline group. This suggests that these pedestrians took large steps to avoid collisions immediately before meeting oncoming neighbors because they were not able to anticipate their neighbors' movements. (*C*) Turn intensity, calculated as the arc length of each step (i.e., product of step length and angle). Error bars are 95% confidence intervals. When compared to the baseline, the values in the other groups are larger, suggesting that pedestrians under the baseline condition did not perform sudden large steps along the *y*-axis (perpendicular to the corridor length), but members of the other groups were forced to perform them, suggesting a difference in their ability to anticipate their neighbors' movements.

Figure S4

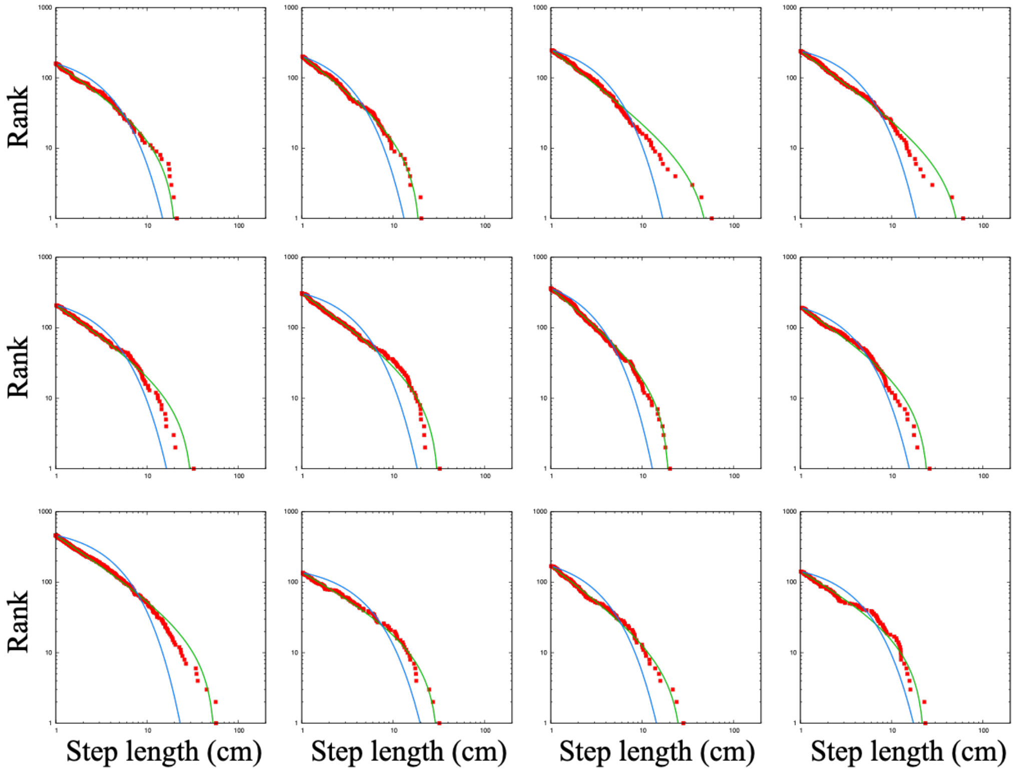


**Fig. S4.** Step-length rank-distribution of the baseline group. Each panel represents a trial. The green and blue lines are the model fits to the truncated power law and exponential distributions, respectively.

Figure S5

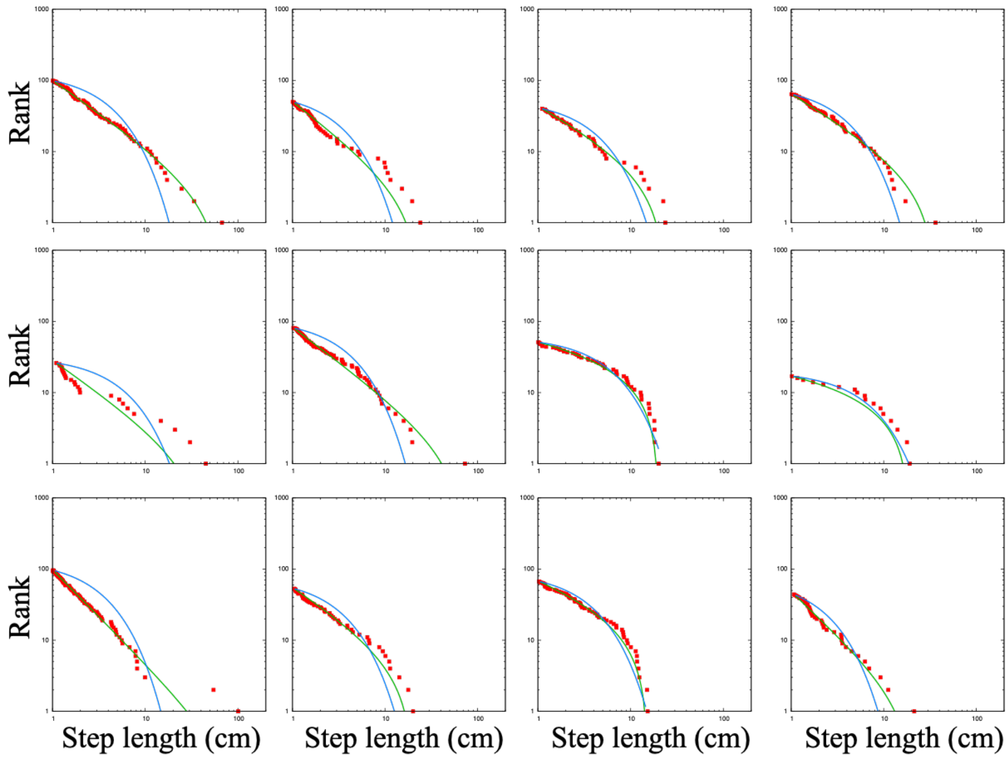


**Fig. S5.** Step-length rank-distribution of distracted pedestrians. Each panel represents a trial. The green and blue lines are the model fits to the truncated power law and exponential distributions, respectively.

Figure S6

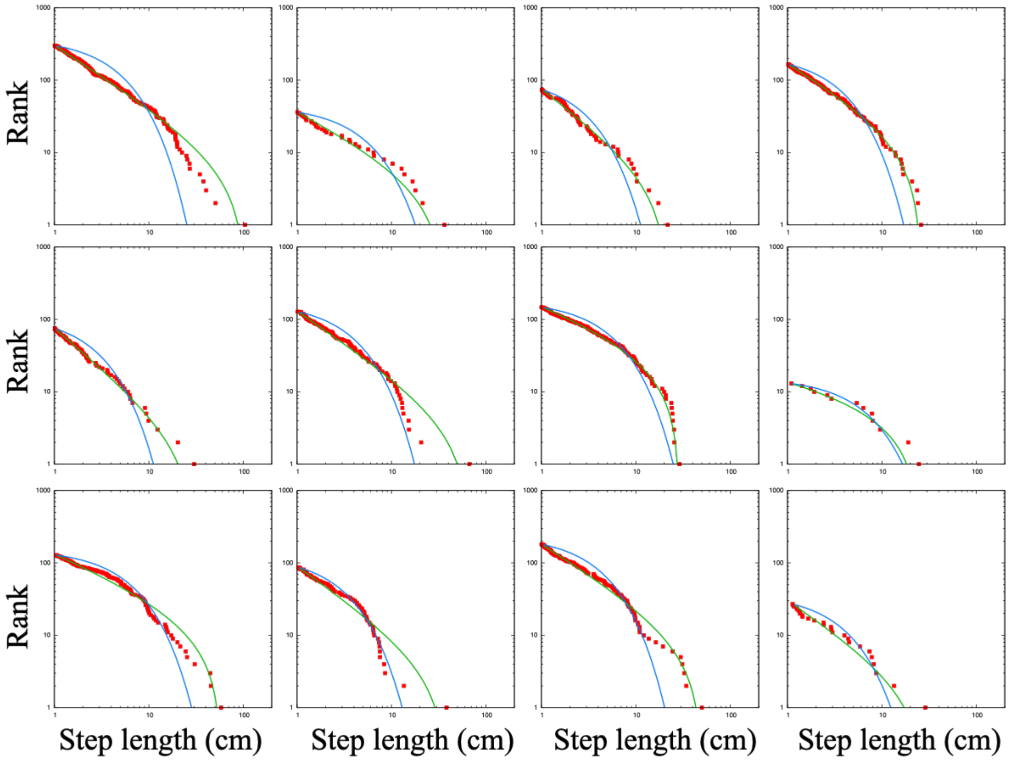


**Fig. S6.** Step-length rank-distribution of pedestrians walking in the same direction as the distracted ones. Each panel represents a trial. The green and blue lines are the model fits to the truncated power law and exponential distributions, respectively.

Figure S7

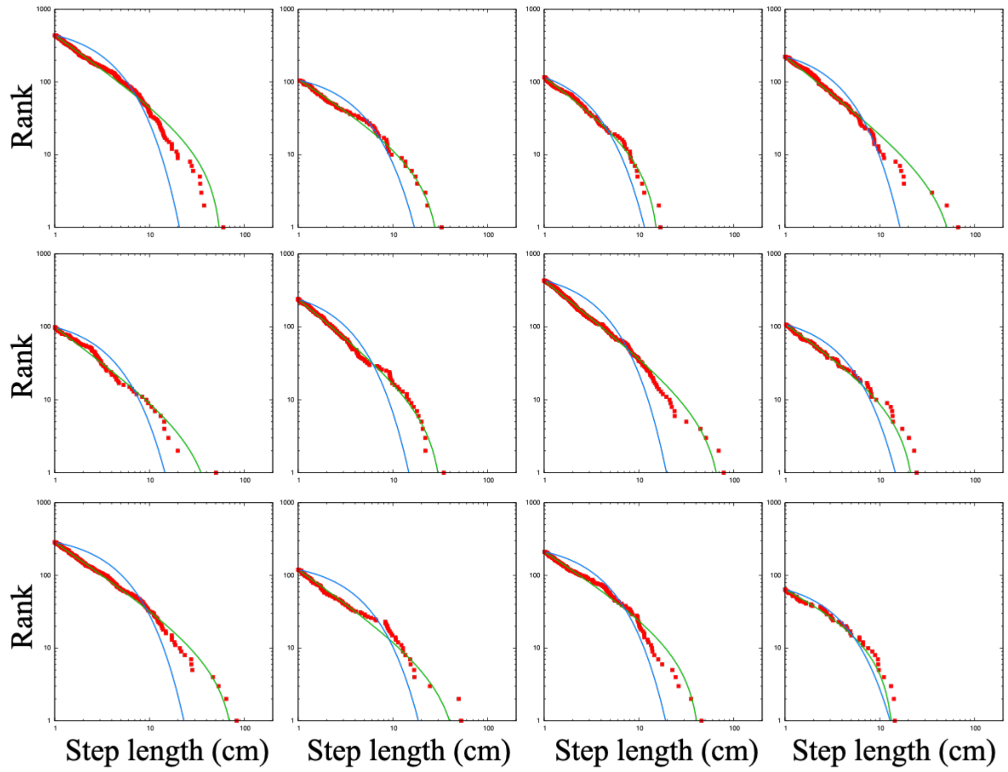


**Fig. S7.** Step-length rank-distribution of pedestrians walking toward the distracted ones. Each panel represents a trial. The green and blue lines are the model fits to the truncated power law and exponential distributions, respectively.

Figure S8

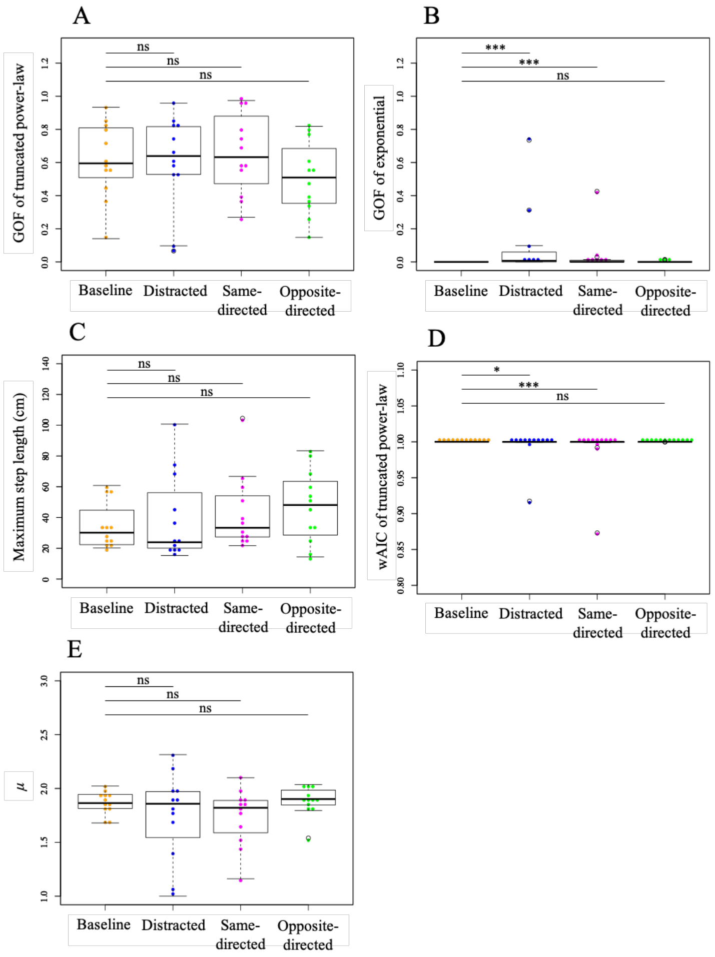


**Fig. S8.** Detailed analysis of (*A*) GOF of the truncated power law, (*B*) GOF of the exponential distribution, (*C*) maximum step length, (*D*) AIC weights of the truncated power law, and (*E*) $\mu$. Each data point represents a trial. Box-and-whisker plots represent the median of the data (central thick line), data between the first and third quartiles (box), data within 1.5× the interquartile range of the median (whiskers), and outliers (unfilled circle). We compared the differences of these values on the distracted, same-directed, and opposite-directed groups from the baseline using Mann–Whitney–Wilcoxon tests with false discovery rate control (*$P$ < 0.05; **$P$ < 0.01; ***$P$ < 0.001; ns, not significant). We found that the GOFs of the exponential distribution of the distracted and same-directed groups were significantly larger than those of the baseline group and that the AIC weights of the truncated power law of the distracted and same-directed groups were significantly smaller than those of the baseline group, suggesting a worse fit of the step-length distribution to the truncated power-law model, as compared with the baseline group.

Figure S9

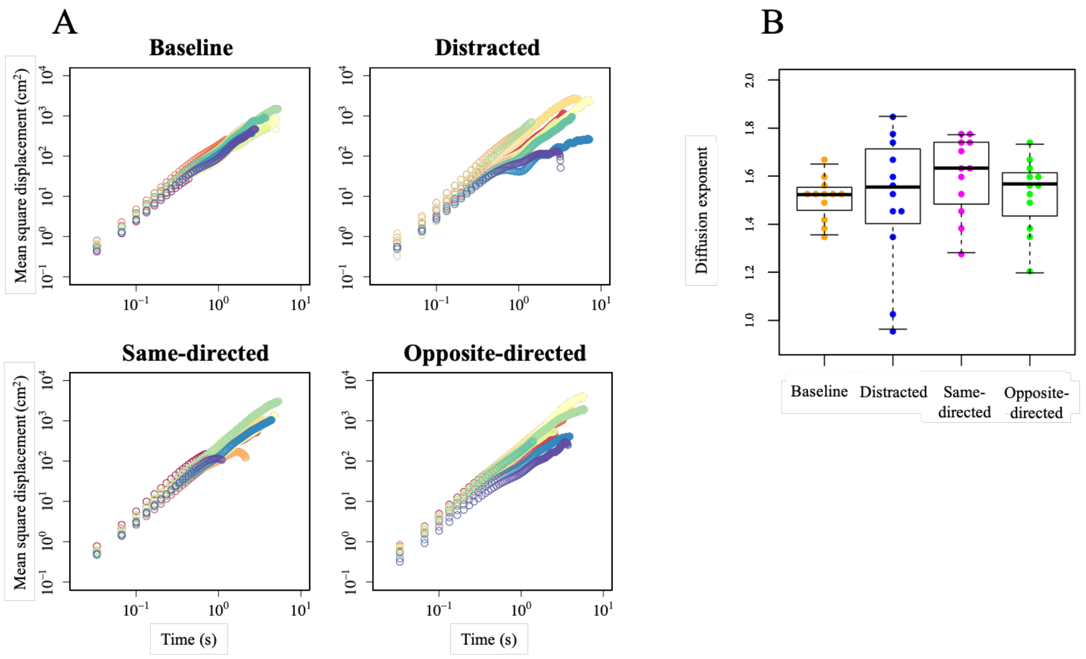


**Fig. S9.** Mean-squared displacement (MSD) along the *y*-axis. (*A*) MSD over time for the baseline and each subgroup under the front condition. Different colors represent different trials. (*B*) Diffusion exponent of the baseline and each subgroup under the front condition. Each data point represents a trial. See the legend for Fig. S8 for a description of the box-and-whisker plots.

Figure S10

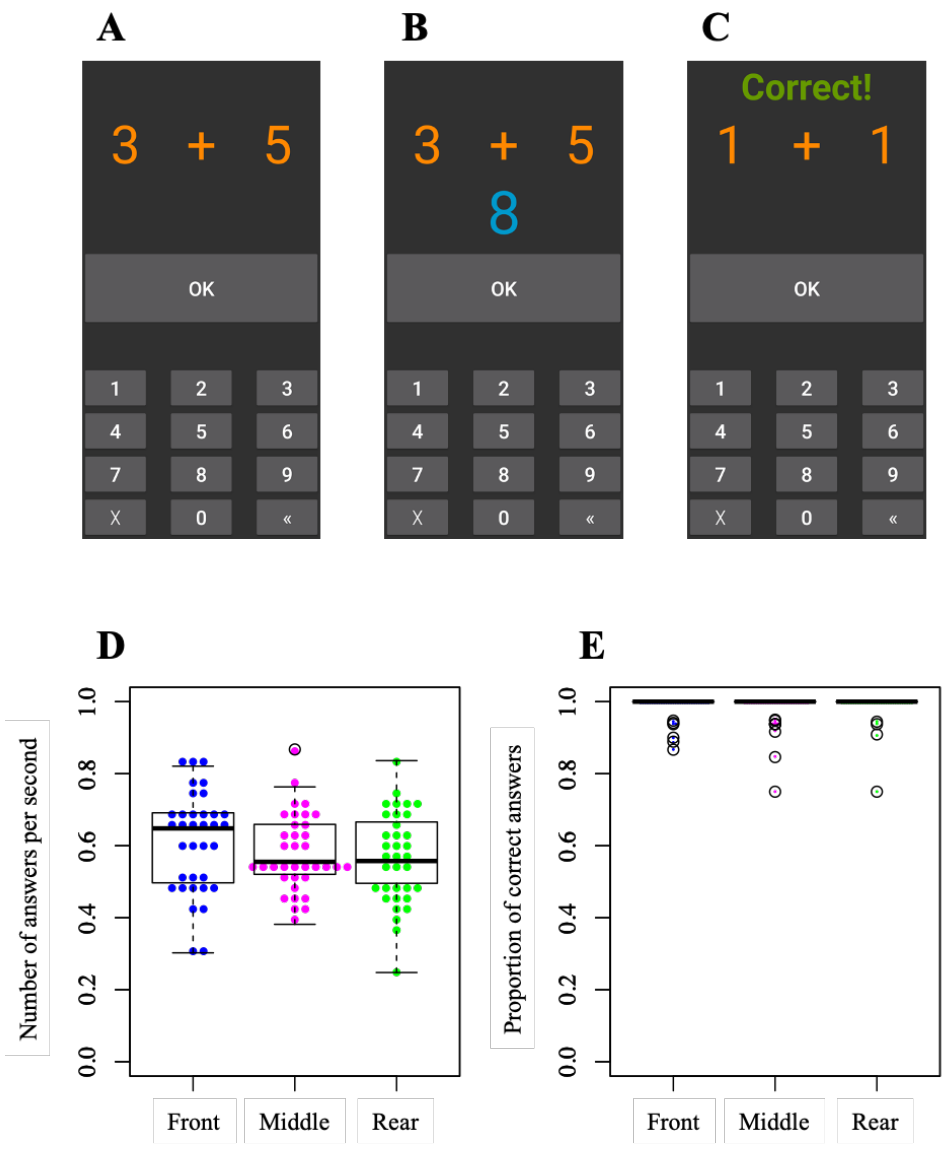


**Fig. S10**. Mobile phone task. (*A*-*C*) Images of the mobile phone task. (*A*) A single-digit addition problem, numeric keyboard, and OK button were initially presented on the display. (*B*) The display after an answer was entered and the OK button was pressed, and (*C*) the next problem, which is presented regardless of whether the previous answer was correct or not. (*D*) Number of answers per second under each condition. (*E*) Proportion of correct answers under each condition. Each data point in (*D* and *E*) represents a distracted pedestrian. See the legend for Fig. S8 for a description of the box-and-whisker plots in (*D* and *E*).

Figure S11

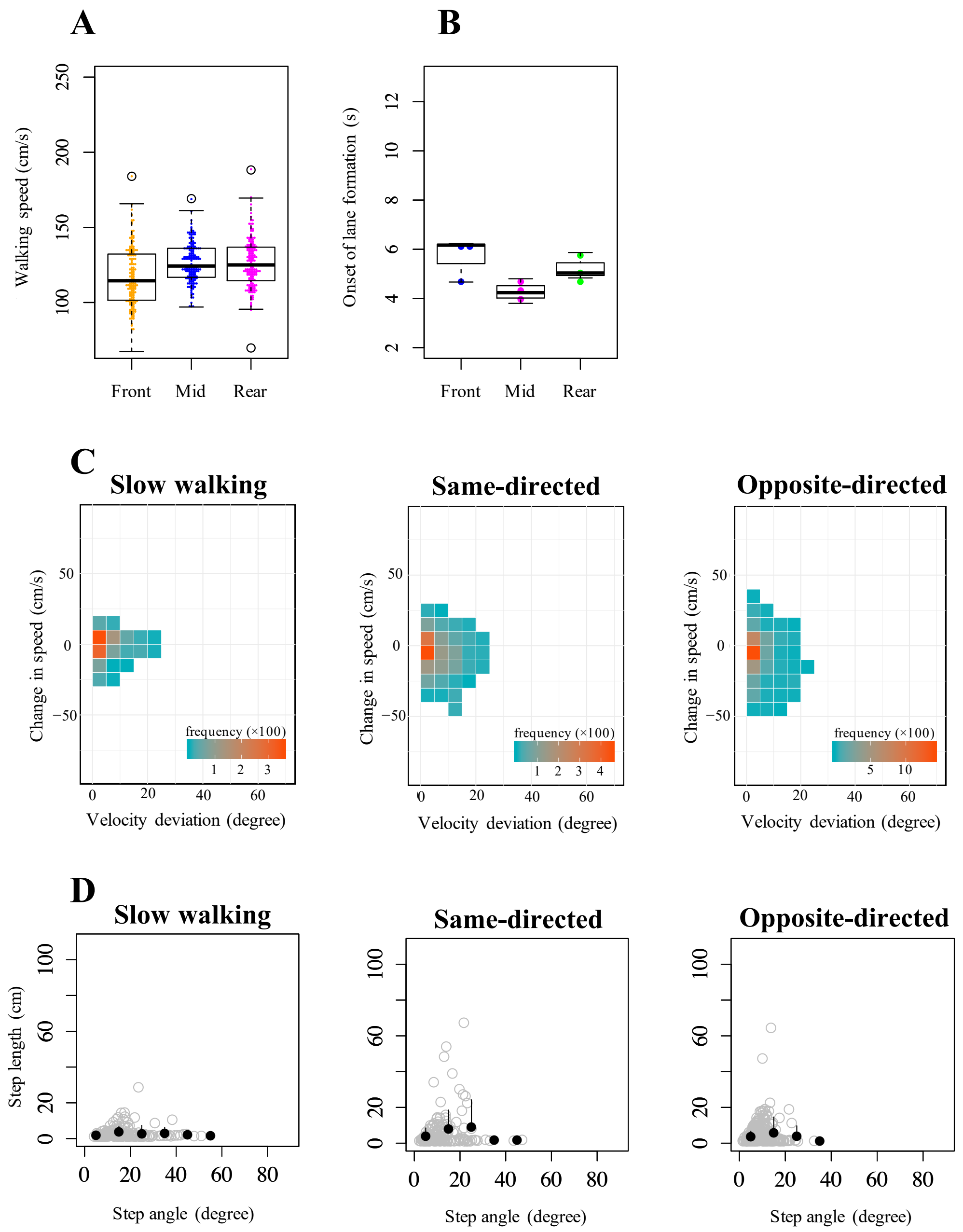


**Fig. S11.** Results of the additional experiments with slow-walking pedestrians. (*A*) Walking speed. (*B*) Onset of lane formation. (*C*) Frequency distribution function of change in speed and velocity deviation with bins of 10 (degrees) × 5 (cm/s) (see also Fig. 3 in the main text). (*D*) Plots of step length against step angle. Black filled circles (error bar: standard deviation) are the binned scattered data (gray circles), with an interval of 10 degrees (see also Fig. S3). See the legend for Fig. S8 for a description of the box-and-whisker plots in (*A* and *B*).
